# Decoding words during sentence production: Syntactic role encoding and structure-dependent dynamics revealed by ECoG

**DOI:** 10.1101/2024.10.30.621177

**Authors:** Adam M. Morgan, Orrin Devinsky, Werner K. Doyle, Patricia Dugan, Daniel Friedman, Adeen Flinker

## Abstract

Sentence production is the uniquely human ability to transform complex thoughts into strings of words. Despite the importance of this process, language production research has primarily focused on single words. However, it remains a largely untested assumption that the principles of word production generalize to more naturalistic utterances like sentences. Here, we investigate this using high-resolution neurosurgical recordings (ECoG) and an overt production experiment where patients produced six words in isolation (picture naming) and in sentences (scene description). We trained machine learning classifiers to identify the unique brain activity patterns for each word during picture naming, and used these patterns to decode which words patients were processing while they produced sentences. Our findings confirm that words share cortical representations across tasks, but reveal a division of labor within the language network. In sensorimotor cortex, words were consistently activated in the order in which they were said in the sentence. However, in inferior and middle frontal gyri (IFG and MFG), the order in which words were processed depended on the syntactic structure of the sentence. Deeper analysis of this pattern revealed a spatial code for representing a word’s position in the sentence, with subjects selectively encoded in IFG and objects in MFG. Finally, we argue that the processes we observe in prefrontal cortex may impose a subtle pressure on language evolution, explaining why nearly all the world’s languages position subjects before objects.

## 1 Introduction

Many animals use symbolic forms of communication: dolphins have names [1], bees dance to signal nectar locations [2], and monkeys and birds use predator-specific calls [3, 4]. While human word (or *lexical*) knowledge is particularly vast, involving tens of thousands of words, the truly remarkable feature of human language is our ability to combine words into sentences, enabling us to express a limitless number of thoughts and ideas.

This communicative ability is central to who we are, but remains poorly understood at the neural level. In particular, the neuroscience of sentence production has been hindered by limitations of traditional noninvasive neural measures, which limit spatial or temporal resolution and are susceptible to motor artifacts, and by the difficulty of experimentally controlling what sentences participants say. Due largely to these challenges, language production research has remained primarily focused on single words, typically employing *picture naming* paradigms where a participant sees a picture of, e.g., a dog, and says “dog.” However, behavioral studies have demonstrated that sentence production is not simply a sequence of single word production tasks [5, 6]. The historical focus on words has left a critical gap in understanding how the brain produces more complex linguistic constructions like sentences.

Among the important insights from research at the word level is that words are not unitary representations. Instead, lexical knowledge involves distinct representations of a word’s semantic (i.e., meaning) [7–9], phonological [8–11], articulatory [12], and grammatical features (*lemmas*) [10, 13]. Each of these representations is associated with distinct cortical regions [14, 15], with articulatory planning in inferior frontal gyrus (IFG) [16, 17]; articulation in sensorimotor cortex (SMC) [18]; feedback in superior temporal gyrus [19] (and visual cortices for sign languages [20]); grammatical features in middle temporal lobe (MTL) [21], and semantics distributed bilaterally throughout cortex [22]. Early production models held that these representational “stages” come online in a strictly feedforward sequence, starting with meaning and ending with articulation (and perception from sensory feedback) [23–25]. However, these models struggled to explain a range of phenomena like speech errors stemming from phonological similarity (e.g., saying *rat* instead of *cat*). Subsequent models therefore introduced an interactive architecture, allowing for bidirectional activation between stages of representation [26–28]. More recently, experimental work has revealed that semantic and phonological information come online at roughly the same time [29, 30], calling into question the notion of stages and leading to the development of models where representations come online in parallel [31, 32].

In contrast to the wealth of knowledge the field has accumulated about single word production, relatively little is known about the type of speech that is unique to our species: sentences [33–35]. Increasingly, researchers are overcoming the obstacles to studying sentence production with non-invasive neural measures [e.g., 17, 33–47]. Many of these studies have focused on the relationship between language production and comprehension, revealing extensive overlap in frontal, temporal, and parietal areas, with additional recruitment of domain-general control regions for production [33, 39, 40, 43, 47] (see [48] for a meta-analysis). Others have examined combinatorial processing [10, 35, 47, 49, 50], revealing that sentence production engages more of the language network – and to a greater extent – than word production [47]. This research has begun to converge on left anterior temporal lobe (ATL) as a hub for composition [50–52], left posterior temporal lobe (PTL) as encoding hierarchical syntax [35, 53, 54] and guiding lexical selection [55–57], and left inferior frontal gyrus (IFG) as responsible for coordinating complex linguistic representations to generate a motor code [10, 16, 35]. Despite these advances, few studies have aimed to directly leverage our understanding of word production to study sentence production. One pressing question, then, is the extent to which the principles of word production generalize to sentences. Here, we address this gap, scaling up from words to sentences by testing the assumption that the mechanisms underlying single word production generalize to more complex linguistic constructs.

To do so, we identify the unique cortical activity patterns that encode six particular words during a picture naming task. We then ask whether these cortical representations are the same for words in list and sentence contexts. By recording electrical potentials directly from the cortical surface in ten neurosurgical patients (ECoG), we achieve high spatial and temporal resolution and avoid motor artifacts, bypassing limitations of traditional non-invasive neural measures. We employ a controlled production experiment (Fig. 1A) and sophisticated machine learning techniques, and demonstrate that words’ cortical representations are in fact shared across tasks. However, the order in which these representations are processed by the brain differs depending on the syntactic structure of the sentence, revealing a dynamic interplay between syntax and word production processes.

**Fig. 1.**
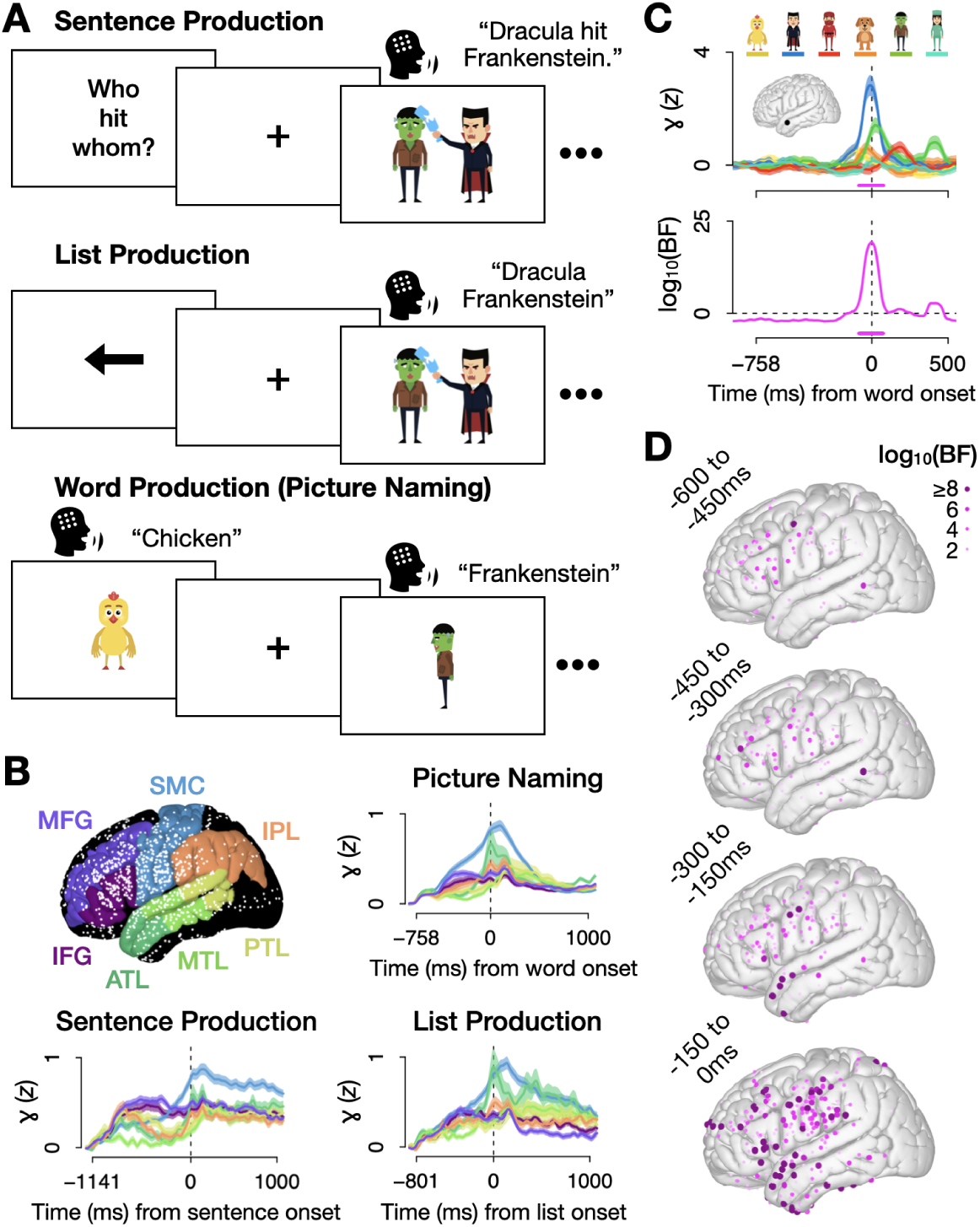
(A) Task design: In sentence production trials, participants described static cartoon scenes in response to preceding questions. Scenes involved two of the six characters used throughout the experiment (*chicken, dog, Dracula, Frankenstein, ninja, nurse*). Half of the questions were manipulated to appear in active syntax (e.g., “Who hit whom?”), implicitly priming active responses (“Dracula hit Frankenstein.”). The other half had passive syntax (“Who was hit by whom?”), priming passive responses (“Frankenstein was hit by Dracula”). In list production trials, participants saw an arrow pointing to the left, in which case they listed the two characters in the subsequent scene from right to left (“Dracula Frankenstein”), or to the right (“Frankenstein Dracula”). In picture naming trials, the six characters repeatedly appeared one at a time and participants responded with a word (e.g., “chicken”). (B) We recorded electrical potentials from 1256 electrodes (white dots) placed directly on the cortical surface in 10 patients. We identified 7 regions of interest (ROIs) in the word production literature. Line plots show the mean neural activity (*z*-scored high gamma amplitude) and standard error per task and ROI, locked to speech onset. (C) A sample electrode: mean activity per word during picture naming (top) and the amount of evidence (BF = Bayes Factor) for word-specific information throughout picture naming trials (bottom; calculated with Bayesian ANOVAs where positive values indicate more evidence for word-specificity). (D) The maximum amount of evidence for word specificity during picture naming in each electrode in the four 150 ms windows leading up to speech onset (*t* = 0).

## 2 Results

We analyzed an existing dataset (first reported in [58]), in which ECoG was recorded from ten neurosurgical patients with electrodes implanted in left peri-Sylvian cortex (Fig. 1B). Patients performed an overt speech production experiment involving the production of the same six words in three tasks: picture naming, list production, and sentence production. In picture naming trials, patients repeatedly saw and named six cartoon characters one at a time. To maximize discriminability, these characters – *chicken*, *dog*, *Dracula*, *Frankenstein*, *ninja*, and *nurse* – differed along a number of dimensions (phonology, number of syllables, proper vs. common noun; see Supplementary Fig. S1). During sentence production, patients overtly described cartoon vignettes depicting transitive actions (e.g., poke, scare, etc.) in response to a preceding question. Questions were constructed using either active syntax (“Who poked whom?”) or passive syntax (“Who was poked by whom?”), implicitly priming patients to respond with the same structure (“The chicken poked Dracula” or “Dracula was poked by the chicken”) [59, 60]. Finally, patients completed a list production task, where the same vignettes as in the sentence production trials were preceded by an arrow rather than a question, indicating the direction in which participants should list the two characters in the scene: left-to-right (e.g., “chicken Dracula”) or right-to-left (“Dracula chicken”). We quantified neural activity as high gamma broadband activity (70 *−* 150 Hz), normalized (*z*-scored) to each trial’s 200 ms pre-stimulus baseline, which correlates with underlying neuronal spiking and BOLD signal [61, 62].

We began by looking at the mean neural activity for each task in seven regions of interest (ROIs) which have previously been implicated in word production [14, 15]. Prior to this and subsequent analyses, we followed previous work [58, 63–67] and temporally warped all trials, setting response times to the median trial duration for each task (-758 ms for picture naming, -1141 ms for sentence production, and -801 ms for list production; see Methods and Supplementary Fig. S2). This boosts signal to noise ratio [58, 66] and tempers extraneous differences. ROIs showed a variety of temporal patterns (Fig. 1B), with the highest levels of activity across tasks achieved in sensorimotor cortex (SMC) during articulation. However, not all of this activity reflects word processing, as various general systems like attention and working memory are also involved in speech production. Notably, many electrodes showed distinct temporal profiles for certain words (Fig. 1C, top). We quantified the amount of evidence for word specificity in each electrode using Bayesian ANOVAs (Fig. 1C, bottom). This information was broadly distributed across cortex, increasing from stimulus onset to speech onset (Fig. 1D; see Supplementary Fig. S3 for each word’s unique network).

To more accurately identify word-specific activity patterns, we performed a series of decoding analyses [68, 69] on the picture naming data. In essence, this analysis (schematized in Fig. 2A) learns the unique pattern of activity for each word in a “training” subset of picture naming trials (90% of trials). Then, it predicts which of the six words a patient is saying in each of the withheld “test” trials (10%) by assessing how similar that trial’s activity pattern is to each of the six learned patterns. If a test trial’s activity pattern is most similar to, e.g., the “chicken” pattern, the classifier will predict that the patient said “chicken.” This would be scored a 1 if the patient was in fact saying “chicken,” and 0 otherwise. By repeating this process for every time sample in every trial, and for 30 combinations of train and test data parcellations (i.e., 10-fold cross validation repeated three times; see Methods), we were able to calculate the mean prediction accuracy over time.

**Fig. 2.**
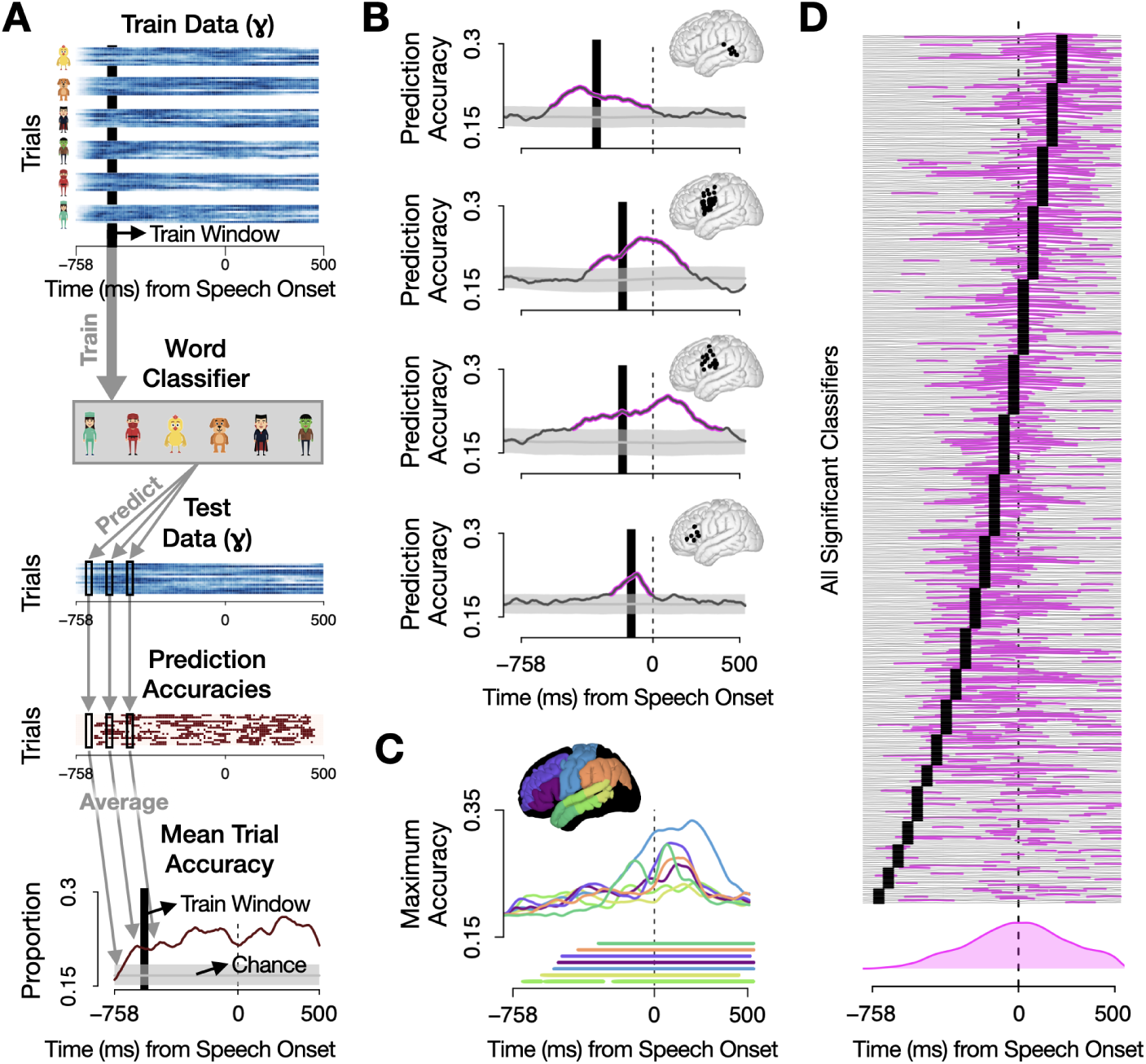
Word-specific information in picture naming. (A) Schematic of the analysis pipeline with simulated data. For each patient and region of interest (ROI), we trained a classifier on word identity using the time-averaged picture naming data from a 50 ms-window (-600 to -550 ms shown in the example; black rectangle). We then predicted word identity in “test” trials that were not in the training data, coding predictions as correct (1) or incorrect (0) for each test trial (row) at each time sample (column), generating a matrix of prediction accuracies (maroon). We repeated this for 30 train/test data parcellations (i.e., 10-fold validation, repeated 3 times), and averaged the resulting prediction accuracies across trials to calculate accuracy over time (bottom). A permutation analysis generated 1000 results reflecting chance performance, and significance was determined with respect to this distribution (gray). This whole process was performed for each patient, each ROI for which a patient had coverage, and each of the 20 training windows spanning -750 to 250 ms, resulting in a total of 1280 classifiers. (B) Prediction accuracies for four sample classifiers with significant predictions (pink highlights; *p <* .05 for 100 ms or *p <* .01 for 50 ms). From top to bottom, training and test data came from (1) Patient 9, PTL, -350 to -300 ms; (2) Patient 5, SMC, -200 to -150 ms; (3) Patient 6, SMC, -200 to -150 ms; and (4) Patient 5, IFG, -150 to -100 ms. (C) Each ROI’s maximum prediction accuracy across classifiers; bars denote significance (*p <* .05 for 100 ms or *p <* .01 for 50 ms). (See Supplementary Fig. S5 for results by ROI.) (D) Prediction accuracies from the 444 classifiers that made significant predictions, stacked vertically to highlight the timecourse of word-specific information in picture naming. Each horizontal line corresponds to one classifier. Pink denotes where that classifier’s accuracy was above chance. Black bars represent time window of training. The pink curve at the bottom shows the density of significant predictions, revealing that the most significant decoding occurred at speech onset (*t* = 0).

We repeated this analysis for each patient and for each ROI. Additionally, because words pass through distinct representational stages (e.g., conceptual, phonological, etc.) [14, 15, 23, 25], the time frame of the training data was consequential. For instance, if we trained the model on just data from *t* = 0 (speech onset), we would likely detect articulatory information and miss semantic, phonological, and grammatical aspects of lexical representation. To avoid this, we spanned time, training models on the mean high gamma activity in each 50 ms window from -750ms to 250ms relative to speech onset. In all, this resulted in 1280 classifiers: 10 patients *×* 7 ROIs *×* 20 training time windows (minus ROIs where patients had insufficient electrode coverage).

Figure 2B shows the prediction accuracies from four sample classifiers. These classifiers predict word identity above chance both before and after their respective training windows (black bars), revealing that whatever information our classifiers encode comes and stays online for longer than 50 ms. In Fig. 2C we plot the maximum accuracy of all classifiers in each region (across patients and training windows), revealing that word identity can be decoded above chance in all seven ROIs (see Supplementary Fig. S4 for all results by ROI). In Fig. 2D, we stacked the time series (like those in Panel B) from all 444 classifiers with significant prediction accuracies (pink highlights). This revealed several patterns. First, the closer a training window (black bar) was to articulation (starting at time 0), the better the decoding, possibly reflecting higher signal-to-noise ratio for articulatory representations than for earlier stages. Second, training times (black) and significant prediction times (pink) tended to overlap, suggesting that the timecourse of representational stages is relatively consistent across picture naming trials. Finally, almost regardless of training time, most classifiers were able to decode above chance at speech onset (*t* = 0). This suggests that pre-articulatory representations stay online at least until speech onset, and post-articulatory representations are engaged throughout production.

To assess whether words have the same cortical representations in picture naming and list production, we tested the generalizability of the picture naming classifiers. We followed the same analysis pipeline depicted in Fig. 2A. However, instead of training and testing on subsets of the picture naming data, we used all of the picture naming data to train classifiers, and then used these classifiers to predict word identity during the production of lists like “Dracula Frankenstein.” The plots in Figure 3A demonstrate results from a sample region, sensorimotor cortex, showing the proportion of trials where SMC classifiers predicted the first word (left; “Dracula,” in our example) and the second word (right; “Frankenstein”). Accuracies are time-locked to the onset of the first word in the left panel and the second word in the right. These sample results come from classifiers trained on data from SMC between 50 and 100 ms after speech onset during picture naming, and averaged across participants. We accurately predict each word as it is being said – e.g., Dracula when the patient says Dracula and then Frankenstein when the patient says Frankenstein. Prediction accuracies from all 97 classifiers that significantly predicted one or both of the two nouns in the list are stacked in Fig. 3B, revealing similar temporal dynamics as we observed in picture naming. The three temporal patterns we observed in picture naming were largely preserved, though to a lesser degree. First, the closer to speech onset the training data came from, the more significant detections the classifier made. Second, significant prediction times tended to overlap with training times, though this was less true for classifiers trained on pre-articulatory data. Finally, significant prediction times tended to overlap with speech onset, particularly for the second word in lists, where many classifiers showed above-chance accuracy during articulation even when they failed to do so during their own training time. Overall, these similarities to the picture naming results (Fig. 2) suggest that list production may involve similar processes as picture naming.

**Fig. 3.**
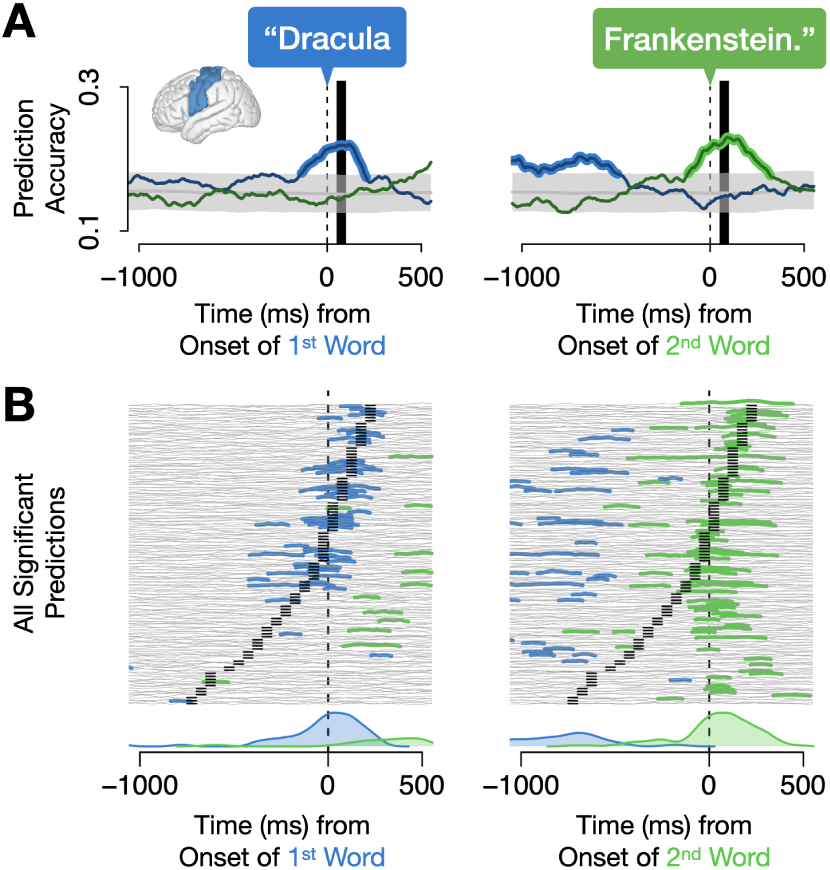
Predicting words in lists using picture naming data. (A) Sample prediction accuracies for the first and second nouns in lists (e.g., “Dracula” and “Frankenstein” in the example) from classifiers trained on picture naming and tested on lists. The significant detections of the two nouns in the list are evidence of words’ common cortical representations across tasks. Training data came from electrodes in SMC (highlighted on brain) between 50 and 100 ms post-speech onset (denoted by black bar) and the resulting prediction accuracies were averaged across patients. Significant prediction accuracy is highlighted in blue for the first word and green for the second (*p <* .05 for 100 ms or *p <* .01 for 50 ms). (B) Prediction accuracy time series (like those in Panel A) from the 97 significant classifiers for lists, stacked vertically to highlight temporal patterns in word-specific information during list production. In this and subsequent stacks of prediction accuracies, each time series (horizontal line) corresponds to the same classifier across the left and right panels. Blue highlights show where a classifier predicted the first noun above chance; green for the second noun (*p <* .05 for 100 ms or *p <* .01 for 50 ms). Black bars denote the time window the training (picture naming) data came from. Blue and green density plots at the bottom summarize the significant predictions. As with picture naming decoding (Fig. 2D), the most significant detections of each word happened at that word’s articulation onset.

There is reason to believe that sentences may behave differently. Whereas word order in lists is linearly structured, word order in sentences is determined by words’ syntactic position in a hierarchical structure, which is in turn based on a complex event-semantic representation. It stands to reason that sentence production may involve fundamentally different mechanisms for accessing and producing words. To test this, we used the same classifiers we trained for the list production analysis to predict word identity during sentences. Like lists, each sentence contained two nouns: the subject and the object, and we recorded the proportion of trials where classifiers predicted each of these. We started by analyzing sentences with active syntax, e.g., “Dracula is hitting Frankenstein,” which, relative to non-canonical structures like the passive, are easier to process and better preserved in aphasic patients [45, 70–73]. Figure 4A shows the results from the same sensorimotor classifiers previously shown for lists in Fig. 3A. For active sentences, subjects and objects were decoded while each was being said (i.e., Dracula and Frankenstein in the sample sentence, but the particular words in subject/object position varied across trials). Stacking the prediction accuracy time series from all 83 significant classifiers (Fig. 4C) revealed that this was a general pattern: subjects and objects were predicted at their respective production times.

**Fig. 4.**
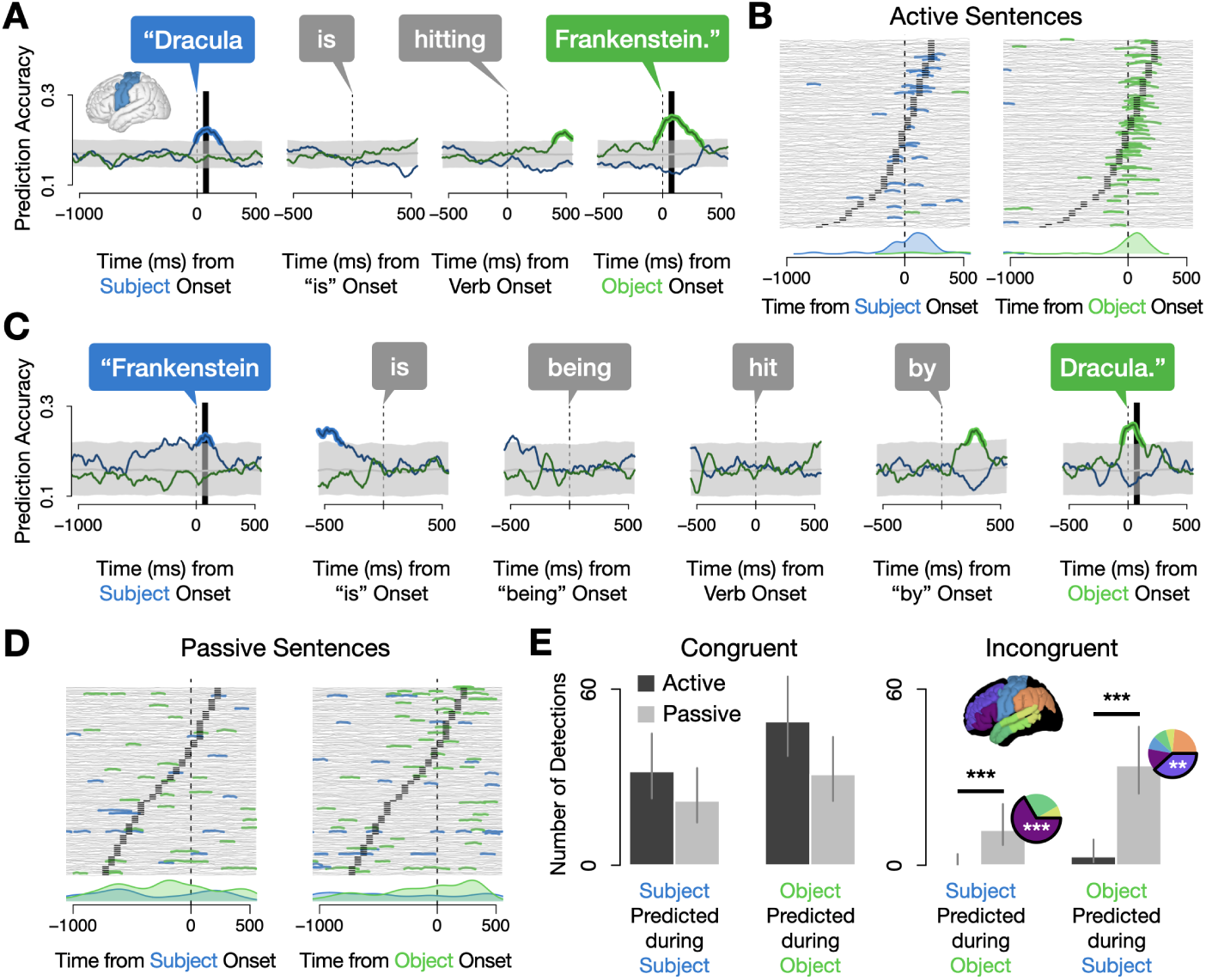
Predicting words in sentences with picture naming data. (A) Sample prediction accuracies for the first and second nouns (i.e., subject and object) of active sentences, locked to the onset of each word. This was the same classifier as in Fig. 3A, i.e., trained on picture naming data from electrodes in SMC between 50 and 100 ms (black bar) after speech onset (averaged across patients). Both nouns were predicted above chance at the time of their respective articulations (blue/green highlights; *p <* .05 for 100 ms or *p <* .01 for 50 ms). (B) Stacked prediction accuracies from the 83 significant classifiers for active sentences, locked to the onset of both nouns. Density plots at bottom show that each word’s accuracy peaked during its articulation. (See Supplementary Fig. S5 for density plots broken down by ROI.) (C) Mean prediction accuracies for passive sentences from the same SMC classifier in Panel A and Fig. 3A. Again, prediction accuracy was above chance for both nouns during their respective articulations. (D) Stacked prediction accuracies from the 97 significant classifiers for passive sentences, locked to each noun’s onset. Unlike in active sentences and lists, there is little correspondence between training time (black bars) and when words were detected (green and blue segments). This point is made especially clear by the density plots, which reveal both subjects and objects were active throughout passive sentence production. (E) Number of classifiers that significantly predicted either word in the sentence, broken down by whether the prediction revealed temporally congruent processing (i.e., detection of the subject during production of the subject or detection of the object during the object) or incongruent (detection of the object during the subject or vice versa). While both active (black) and passive (gray) sentences involved temporally congruent word processing, passives had significantly more incongruent detections than actives for both incongruent subjects (*χ*^2^(1) = 10.132, FDR-corrected *p <* .001; here and for remaining tests in this caption: one-sided Test of Equal Proportions) and incongruent objects (*χ*^2^(1) = 24.687, FDR-corrected *p <* .001). Pie charts show where these incongruent detections were made: incongruent passive subjects were detected in IFG more than any other ROI (*χ*^2^(1) = 23, FDR-corrected *p <* .001) whereas incongruent passive objects were found in MFG (*χ*^2^(1) = 11.421, FDR-corrected *p <* .001; see Supplementary Fig. S5 for more detail).

However, in active sentences, the order of the two nouns is confounded with their relative salience in the event. That is, the character performing the action is the subject and comes first, and the character being acted upon is the object and comes second. It is therefore not entirely surprising that the brain processes words in active sentences the way it does in lists: in the order that they are produced. We wondered whether this pattern was generally true of sentence production, or only true when word order aligns with salience. A potentially interesting test case is the English passive (e.g., “Frankenstein is being hit by Dracula”), which involves reversing the order of nouns – i.e., producing the character being acted upon first, and the character doing the action second (see [45] for discussion of the neural processing of canonical vs. noncanonical word orders).^1^ Passive sentences thus present an opportunity to disentangle serial order and salience, as they convey the same meaning but reverse the order of words. We expected that in sensorimotor cortex, where articulatory information is encoded, words would be decoded in the same order as in speech. Figure 4C shows the predictions of the same SMC classifier from Panel A (and Fig. 3A) for passive sentences, and the results did in fact show this temporal congruence. To look at the overall pattern, we stacked the predictions from all 97 classifiers that made above-chance predictions (Fig. 4D; for a breakdown by ROI see Supplementary Fig. S5). This analysis revealed an entirely different pattern of results. In passives, both the subject and object remained active throughout the entirety of the sentence, revealing that the brain processes both characters in the sentence simultaneously rather than sequentially. To assess whether this constituted a statistically significant difference from active sentences, we counted the number of classifiers that detected each word during the production of each word (Fig. 4E) – i.e., the number of classifiers that detected the subject when the subject was being said or the object when the object was being said – “congruent” detections – and the number of classifiers that detected the object when the subject was being said or the subject when the object was being said – “incongruent” detections. In both active and passive sentences we observed many congruent detections (Fig. 4E, left). However, of the incongruent detections we observed (Fig. 4E, right), nearly all were in passive trials. Relative to active sentences, passive sentences involved significantly more incongruent detections of both subjects (*χ*^2^(1) = 10.132, FDR-corrected *p <* .001) and objects (*χ*^2^(1) = 24.687, FDR-corrected *p <* .001; both one-sided Tests of Equal Proportions).^2^ This pattern was driven by two regions (see pie charts and Supplementary Fig. S5): IFG, which preferentially encoded subjects throughout passive sentences, and MFG, which preferentially encoded objects. Strikingly, even when patients were producing the subject of a passive sentence, their brains more strongly encoded the object, as evidenced by the higher number of incongruent detections (predictions of the object) than congruent ones (predictions of the subject), although this difference was not statistically significant after corrections for multiple comparisons. In summary, our decoding analysis revealed a marked difference in lexical processing between active and passive sentences. While actives exhibit a pattern similar to that in list and single word production, where words are activated in the order they are produced, passive sentences involve sustained, simultaneous encoding of both the subject and object nouns. This difference was region-specific, with sensorimotor cortex consistently representing lexical information in task-agnostic ways, but IFG and MFG showing sensitivity to syntactic structure, highlighting a division of labor within the language network where prefrontal regions support structure-dependent processing demands.

## 3 Discussion

Single word production tasks like picture naming have dominated the neuroscience of language production. In this study, we leveraged the unparalleled spatiotemporal precision of ECoG and employed an innovative cross-task classification approach to demonstrate similarities and differences in word processing between single word production and sentence production. We first demonstrated that individual words can be decoded from patterns of activity in picture naming data, verifying that our data contained word-specific information. Next, we used the picture naming data to train a series of classifiers on word identity at 20 time points over the course of picture naming and in seven regions of interest. We showed that these classifiers successfully decoded each word during the production of lists of words in the order in which words were said. We then applied these same classifiers to sentences to determine whether sentence production could similarly be modeled as a sequence of single word productions. We started with active sentences, which represent the canonical word order in English and typically involve producing nouns in order of agency, from most agentive to least. As we observed for lists, classifiers successfully decoded the two nouns in active sentences in the order in which they were produced. Finally, we used the classifiers to decode word identity during the production of passive sentences, which encode the same events as active sentences but with the reverse word order. Here, we observed a significant departure from the temporal alignment of word processing in the brain and word order in speech: rather than encoding each word as it was being said, the brain encoded both words simultaneously for the duration of the sentence. Furthermore, our data revealed a spatial code in prefrontal cortex for words’ structural position, with subjects preferentially encoded in IFG and objects in MFG.

Our findings validate various aspects of cognitive [74–76] and computational [77, 78] models of sentence production. These models assume that word representations are invariant across different behaviors, which is reflected in our finding that the cortical representation of words during picture naming generalizes to both list and sentence contexts. Furthermore, these models build in a dependence on syntactic structure during sentence planning, predicting that the dynamics of word processing may vary with syntactic structure. Consistent with this, we observed striking differences between active and passive sentences: in actives, the subject and object were decoded sequentially in the order in which they were produced, whereas in passives we continuously decoded both nouns throughout the duration of the sentence. This sustained representation was driven by activity in IFG and MFG, aligning with prior experimental [79–81], stimulation [82], and lesion-symptom mapping studies [83, 84] implicating these prefrontal regions in the processing of noncanonical structures like passives.

There is ongoing debate regarding the specific functions of IFG and MFG in passives. While we have casually attributed these differences to “syntax,” they could in principle derive from other differences between actives and passives. Indeed, the literature identifies a number of possibilities including working memory [85], thematic role assignment [80], and syntactic movement – a syntactic operation associated with certain complex structures [79, 81, 83]. However, much of this prior work relies on measures like “activity,” which do not directly map onto specific psychological processes or representations, complicating efforts to discern among these possibilities. By tracking word-specific information, our findings reveal at least one function of IFG and MFG in passives: the sustained encoding of words and their syntactic roles. This finding poses a challenge for the syntactic movement account, as movement pertains to abstract structure rather than specific words [86, 87]. Similarly, the thematic role assignment hypothesis is inconsistent with our results, as thematic roles did not differ between actives and passives in our study.

Instead, our results align most strongly with a working memory (or perhaps attention) account. Passives likely require more engagement of such cognitive resources, as they have lower frequency than actives, more words, and, in our stimuli, involve the reversal of the canonical agent-first ordering. We suspect that this latter property drives the effect in our data given that IFG and MFG encoded words according to their syntactic role in the sentence. Ongoing access to information about which noun occupies which syntactic position may be needed to override default syntactic processes, which would presumably favor mapping agents to subject, as in active sentences. Thus, the sustained encoding of nouns in IFG and MFG may constitute a core neural mechanism by which the brain exerts top-down control of speech, enabling flexible sentence production to meet situational or task demands. A working memory account further aligns with the fact that IFG and MFG sit squarely within the *multiple-demand network*, a domain-general system that supports cognitive resources like attention and working memory [88–90]. While this explanation best fits our findings, it does not rule out the possibility that these regions additionally support other functions like thematic role assignment and syntactic movement. Importantly, the relative contributions of these functions may differ between production and comprehension, meaning that our results may not be directly comparable to other experimental work, which has primarily focused on comprehension.

Syntactic roles, or structural positions like subject and object, are key parts of all models of language processing. They are necessary for mapping words to semantic/thematic roles (i.e., who performs an action vs. who it is done to). In production, these roles must be quickly and flexibly linked to specific words so that, for example, “Dracula” can be the subject of one sentence and the object of another in rapid succession without causing confusion (see discussion of the “fast-changing weights” in Chang et al.’s 2006 computational model). Theoretical accounts often point to coherence as a possible neural mechanism for this type of binding [78, 91], but there remains little empirical evidence for this (or any) particular implementation. Here, we uncovered a spatial code for syntactic roles during passives, with subjects encoded in IFG and objects in MFG.

Two caveats, however, warrant consideration. First, while active sentences also have subjects and objects, we found no evidence for role-specific encoding in pre-frontal cortex during actives (see Supplementary Fig. S5), despite higher statistical power (patients produced approximately three times more actives than passives). This suggests that the spatial code we observed is not the general neural mechanism for encoding syntactic role (or linear position, whichever it may be). However, an alternative explanation is that the brain does not rely on syntactic roles during the production of actives. For instance, speakers may develop a heuristic strategy for producing very common structures [92], for instance, something like “produce the more agentive noun first.” Indeed, previous work has shown that sentence production is not always driven by bottom-up syntactic encoding in production, and can instead be driven by attention [93] (see [75] for discussion). The second caveat is that the absence of evidence for position-specific encoding in actives limits our ability to disentangle whether this feature of passives encodes syntactic roles or linear position. Prior work has linked IFG to linear rather than hierarchical-syntactic representations [54], potentially indicating a linear position interpretation of our results is a better fit. However, further work is needed to conclusively tease apart these possibilities. Regardless, the pattern we observed for passives is to our knowledge the first demonstration of a neural code for a noun’s position in a sentence, be it linear or hierarchical. This finding demonstrates how a spatial organization of lexical information can and does encode positional information in language.

Future work is also needed to identify *which* lexical representations we detected. Our stimuli, designed with different analyses in mind, do not readily distinguish between different types of lexical information. For example, an electrode that selectively responds to “Dracula” and “Frankenstein” might reflect semantic information (*monsterhood*) or form-level information, as both words have stress on the first of three syllables. One approach to isolating lexical stages could involve examining the spatial or temporal distribution of lexical information. For instance, if different lexical representations correspond to specific ROIs and come online in a feedforward sequence during picture naming, then one would expect word-specific information to first emerge in conceptual regions, then in lemma regions, and so on. However, it remains an open question whether either of these assumptions are valid. Spatially, there is growing consensus that at least semantic knowledge is broadly distributed across cortex [22], which, if true, would mean that decoding in any ROI could be driven by semantic information. Indeed, this could be true of other types of lexical representations as well, where evidence is stronger for a more spatially concentrated code, but still not a matter of certainty. Temporally, if parallel models of word production are correct and lexical representations come online contemporaneously [30–32], then temporal patterns are of limited use in distinguishing between representations. But even under strictly feed-forward circumstances, our data reveal another complication: the signal-to-noise ratio of different representations appears to change over time, leading to potentially sizable differences between when a representation comes online and when it is detected. Evidence for this is clearest in the picture naming data (Fig. 1D), where words tended to be decoded around the onset of articulation regardless of when the training data came from. This was particularly true for earlier training times (the lower part of the “stack” plot), where many classifiers successfully decoded words around time 0 even when they were unable to do so at the time they were trained on. This points toward a signal-to-noise ratio increase leading up to articulation, making information easier to decode at articulation even if it came online much earlier. Thus, we caution against directly interpreting the lack of decoding in classifiers trained at earlier times, as false negatives are likely to become more frequent the earlier the training data come from.

Interestingly, there was one clear exception to this trend. Lexical information in posterior temporal lobe (PTL) peaked far earlier than in other ROIs (Supplementary Fig. S4B), consistent with evidence implicating this region in phonological encoding (see [15] for a review). However, other work has associated PTL with more general processes such as lexical access and coordinating different types of information [55–57], which would implicate multiple lexical representations. Thus, while there are suggestive features of our data, ongoing disagreements in the field about the timecourse and localization of lexical representations, combined with limitations of our stimulus design prevent us from answering these questions definitively. However, this study introduces an innovative approach to investigating these issues – one that we anticipate will be instrumental in resolving these questions in future research.

Finally, we suggest that our findings may shed light on a widely noted but poorly understood pattern among the worlds’ 6,000 languages. Specifically, there are six possible ways a language can arrange subjects, verbs, and objects: Subject-Verb-Object (as in the English “I eat cake”), Subject-Object-Verb (as in Farsi “*man keik mikho-ram*,” literally “I cake eat”), and so on. However, of these 6 logically possible word orders, fewer than 5% of languages place objects before subjects (e.g., Object-Subject-Verb) [94, 95]. One possible reason for this is that subjects tend to be more agentive (semantically salient) than objects, and there is a natural tendency in speech to order words from most to least agentive (a bias apparently preserved across hominids [96]). In our experiment, passive sentences provided an opportunity to visualize how the brain processes words when producing less agentive words before more agentive ones. In these cases, word planning involved a much more complex temporal pattern. Indeed, whereas word planning in actives resembled picture naming and lists, in passives the brain encoded both the subject and the object for the duration of the sentence. This was driven by sustained activation of both nouns in prefrontal cortex. Specifically by IFG sustained a representation of the subject, and MFG sustained a representation of the object. Furthermore, reaction times, commonly interpreted as an index of processing difficulty, were significantly longer for passives than both actives and lists (see Section A.6 in Supplementary Information). Taken together, these facts point toward a processing-based explanation of the cross-linguistic dominance of subject-before-object word orders like those in English and Farsi. Producing words in order from least to most salient may simply be harder for the production system. We speculate that, over the course of language evolution, this difficulty exerts a subtle pressure on language change, making it more likely for languages to evolve in the direction of subject-before-object orders.

## 4 Methods

The data in this study were also reported in Morgan et al. (2024). Details of participants, experimental design, and data collection are repeated below.

### 4.1 Participants

We recorded data from ten neurosurgical patients undergoing evaluation for refractory epilepsy (3 female, mean age: 30 years, range: 20 to 45). All ten were implanted with electrocorticographic grids and strips. Patients provided informed consent both in writing and then again orally prior to the beginning of the experiment. The implantation and location of electrodes were guided solely by clinical requirements. Eight participants were implanted with standard clinical electrode grid with 10 mm spaced electrodes (Ad-Tech Medical Instrument, Racine, WI). The remaining two participants consented to a research hybrid grid implant (PMT corporation, Chanassen, MN) that included 64 additional electrodes between the standard clinical contacts (with overall 10 mm spacing and interspersed 5 mm spaced electrodes over select regions), providing denser sampling but with positioning based solely on clinical needs. The research study protocol was approved by the NYU Langone Medical Center Committee on Human Research.

### 4.2 Data collection and preprocessing

Participants were tested while resting in their hospital bed in the NYU Langone epilepsy monitoring unit. Stimuli were presented on a laptop computer screen positioned at a comfortable distance from the participant. Participants’ voice was recorded with a cardioid microphone (Shure MX424). The experiment computer generated inaudible TTL pulses marking the onset of a stimulus. These were recorded in auxiliary channels of both the clinical Neuroworks Quantum Amplifier (Natus Biomedical, Appleton, WI), which records ECoG, and the audio recorder (Zoom H1 Handy Recorder). The microphone signal was also fed to the audio recorder and the ECoG amplifier. These redundant recordings were used to sync the speech, experiment, and neural recordings.

The standard implanted ECoG arrays consisted of 64 macro-contacts (2 mm exposed, 10 mm spacing) in an 8*×*8 grid. Hybrid grids contained 128 electrode channels, including the standard 64 macro-contacts plus 64 additional interspersed smaller electrodes (1 mm exposed) between the macro-contacts (providing 10 mm center-to-center spacing between macro-contacts and 5 mm center-to-center spacing between micro/macro contacts, PMT corporation, Chanassen, MN). The FDA-approved hybrid grids were manufactured for research purposes, which we explained to patients during consent. In all ten patients, ECoG arrays were implanted on the left hemisphere. The location of the grid was solely dictated by clinical needs.

ECoG was recorded at 2,048 Hz, which was decimated to 256 Hz prior to processing and analysis. We excluded electrodes with artifacts (i.e., line noise, poor contact with the cortex, and high amplitude shifts) or with interictal/epileptiform activity prior to subtracting a common average reference (across all valid electrodes and time) from each individual electrode. We then extracted the envelope of the high gamma component (the average of three evenly log-spaced frequency bands from 70 to 150 Hz) from the raw signal with the Hilbert transform.

The signal was epoched locked to stimulus (i.e., cartoon images) and production onsets for each trial. The 200 ms silent period preceding stimulus onset (during which patients were not speaking and fixating on a cross located at the center of the screen) was used as a baseline, and each epoch for each electrode was *z*-scored (i.e., normalized) to this baseline’s mean and standard deviation.

### 4.3 Experimental Design

#### 4.3.1 Procedure

The experiment was performed in a single session that lasted approximately 40 minutes. Stimuli were presented in pseudo-random order using PsychoPy [97]. All stimuli were constructed using the same 6 cartoon characters (chicken, dog, Dracula, Frankenstein, ninja, nurse), chosen to vary along many dimensions (e.g., frequency, phonology, number of syllables, proper vs. common, etc.) to facilitate identification of word-specific information at analysis.

The experiment had a blocked design. Blocks were ordered in terms of importance to ensure that the most valuable data was collected first, in case a patient ceased participation mid-way through (e.g., due to discomfort, fatigue, or seizure activity). The experiment began with two short familiarization blocks. In the first block (6 trials), participants saw each of the six cartoon characters once with labels (*chicken, dog, Dracula, Frankenstein, ninja, nurse*) written beneath the image. Participants read the labels aloud, after which the experimenter pressed a button to go to the next trial. In the second block, participants saw the same six characters one at a time, twice each with order pseudo-randomized (12 trials), but without labels. Participants were instructed to name the characters out loud. After naming the character, the experimenter pressed a button revealing the target name to ensure patients had learned the correct labels. Participants then completed the first picture naming block (96 trials). Characters were again presented in the center of the screen, one at a time, but no labels were provided.

Next, participants performed a sentence production block (60 trials), which began with two practice trials. Participants were instructed that there were no right or wrong answers, that the goal of the experiment was to understand what the brain is doing when people speak naturally. On each trial, participants saw a 1 s fixation cross followed by a written question, which they were instructed to read aloud, ensuring attention. After another 1 s fixation cross, a static cartoon vignette appeared in the center of the screen depicting two of the six characters engaged in a transitive event (one character acting on the other). Participants were instructed to respond to the question with a description of the vignette. The image remained on the screen until the participant completed their response, at which point the experimenter pressed a button to proceed. After the first 12 trials, the target sentence (i.e., an active sentence after an active question or passive sentence after a passive question) appeared in text on the screen and participants read it aloud. We described these target sentences as “the sentence we expected you to say.” The goal of this was to implicitly reinforce the link between the syntax of the question and the target response. If the participant appeared to interpret these as corrections, the experimenter reminded them that there were no right or wrong answers.

Between each sentence production trial, we interleaved two picture naming trials, which was demonstrated to reduce task difficulty and facilitate fluent sentence production during pilot testing. The picture naming trials showed the two characters that would be engaged in the subsequent vignette, presented in a counterbalanced order such that on half of trials they would appear in the same order as in the target sentence response, and in the opposite order on the other half.

After the sentence block, participants performed the listing block. The list production block was designed as a secondary control condition in the original study [58], and as such it was ordered last. List production was designed to parallel sentence production. Each trial began with a 1 s fixation cross, followed by an arrow pointing either left or right appeared for 1 s in the center of the screen. After another 1 s fixation cross, a cartoon vignette, taken from the exact same stimuli as in the sentence block, appeared on the screen. Participants named the two characters in the vignette either from left to right or from right to left, according to the direction of the preceding arrow. As in sentence production trials, each list production trial was preceded by two picture naming trials.

Between each block, participants were offered the opportunity to end the experiment if they did not wish to continue. One participant stopped before the list production block, providing only data for picture naming and sentence production. The remaining nine participants completed all three blocks. These nine were also offered the opportunity to complete another picture naming block and another sentence production block. Six consented to an additional picture naming block and two additionally consented to another sentence production block.

#### 4.3.2 Stimulus Design, Randomization, and Counterbalancing

Picture naming stimuli consisted of images of the 6 characters presented in pseudorandom order so that each consecutive set of 6 trials contained all 6 characters in random order. This ensured a relatively even distribution of characters over time, and that no character appeared more than two times in a row. Characters were pseudorandomly depicted in 8 orientations: facing forward, backward, left, right, and at the 45*^◦^* angle between each of these.

Sentence production stimuli consisted of a written question followed by a static cartoon vignette. Questions were manipulated so half were constructed with passive syntax and the other half with active. All questions had the format: “Who is [verb]-ing whom?” or “Who is being [verb]-ed by whom?”. There were 10 verbs: *burn, hit, hypnotize, measure, poke, scare, scrub, spray, tickle, trip*. Each verb was used to create 3 vignettes involving 3 characters in a counterbalanced fashion so that each character was the agent (i.e., active subject) in one vignette and the non-agent (i.e., active object) in one vignette. Each of these three vignettes was shown twice in the sentence production block, once preceded by an active question and once by a passive question, priming active and passive responses [59, 60]. Vignettes were flipped around the vertical axis the second time they appeared so the character that was on the left in the first appearance was on the right in the second appearance. This was also counterbalanced so that on half of the trials in each syntax condition (active/passive) the subject was on the left. List production stimuli consisted of the same 60 vignettes, also pseudorandomly ordered and counterbalanced across conditions (i.e., arrow direction) so that (a) on half of trials the first character to be named appeared on the left, and (b) on half of trials the first character to be named was the agent of the depicted event.

### 4.4 Data Coding and Inclusion

Speech was manually transcribed and word onset times were manually recorded using Audacity [98] to visualize the waveform and spectrogram of the audio recording. Picture naming trials were excluded if the first word uttered was not the target word (e.g., “Dracula – I mean Frankenstein”). Sentence trials were excluded if the first word was incorrect (i.e., “Dracula” instead of “Frankenstein,” regardless of active/passive structure) or if the meaning of the sentence did not match the meaning of the depicted scene; no sentences were excluded because the syntax did not match that of the prime question. Sentences were coded as active or passive depending on the structure the patient used, not the prime structure. Listing trials were excluded if the first word was incorrect (“Dracula” instead of “Frankenstein”) or if the order did not match that indicated by the arrow.

In analyses for the three trial types (picture naming, sentence production, and list production), data from all patients who completed trials in that block are included. Data from one patient who did not complete the list production block is not included in the list production analyses, and data from 3 patients who produced 3 or fewer passive sentences during sentence production blocks were not included in the analyses of passive sentences.

### 4.5 Electrode Localization

Electrode localization in both subject space and MNI space was based on coregistering a preoperative (no electrodes) and postoperative (with electrodes) structural MRI (in some cases, a postoperative CT was employed depending on clinical requirements) using a rigid-body transformation. Electrodes were then projected to the cortical surface (preoperative segmented surface) to correct for edema-induced shifts following previous procedures [99] (registration to MNI space was based on a nonlinear DARTEL algorithm). Based on the subject’s preoperative MRI, the automated FreeSurfer segmentation (Destrieux) was used for labeling electrodes’ within-subject anatomical locations.

### 4.6 Significance testing and corrections for multiple comparisons in time series data

Statistical tests on time series data were performed independently at each time sample, producing the same number of *p*-values as there are samples in the time series. To correct for multiple comparisons we follow [58, 100, 101] and establish a conservative criterion for significance for all time series comparisons: an uncorrected *p*-value that remains below .05 for at least 100 consecutive milliseconds or below .01 for at least 50 consecutive milliseconds.

### 4.7 Multi-class classification

For the analyses in Figs. 2–4 we trained multi-class classifiers on word identity using the *caret* and *nnet* packages [102, 103] in R [104]. Classifiers consisted of a series of one-vs-rest logistic regressions (fit as a neural network), which were chosen for their simplicity and interpretability. For the picture naming analyses (Fig. 2), we used a repeated cross-fold validation procedure (3 repeats, 10 folds) to calculate prediction accuracy, and arbitrarily chose a mid-range value of 10*^−^*^3^ for decay, the lone hyperparameter in this model (typical values are logarithmically spaced along the range from 10*^−^*^6^ to 10^0^). For the subsequent analyses of list and sentence production data, we first performed repeated cross-validation on the picture naming data again to find the optimal hyperparameters for each individual classifier, and then retrained each model with that hyperparameter and using all of the picture naming trials (rather than a training subset). We then used this model to predict word identity at every time point throughout each trial in the list and sentence production blocks. Prediction accuracy time series like those in Fig. 4A reflect the mean of the binary accuracy scores across sentence production trials separately for the subject and the object (which on different trials were different combinations of the six nouns, e.g., Dracula, dog, etc.), smoothed with a 100 ms boxcar function. To generate the noise distribution (gray shaded area), we performed a permutation analysis, shuffling labels on the test data and repeating the prediction analysis 1000 times. We determined significance by calculating the upper 95th and 99th percentiles of the mean trial accuracies generated by the permutation analysis for each time sample (see Section 4.6 for details on multiple comparisons corrections).

### 4.8 Temporal warping

The time between stimulus and speech onsets the *planning period*, varied both across and within patients. Consequently, cognitive processes become less temporally aligned across trials the farther one moves away from stimulus onset in stimulus-locked epochs or from speech onset in speech-locked epochs. Temporal warping reduces such misalignments [63–67], which we previously verified in this dataset [58]. Following [58, 66], we linearly interpolated the data in the middle of the planning period (from 150 ms post-stimulus to 150 ms pre-speech) for each trial, setting all trials’ planning periods to the same duration (Supplementary Fig. S2): the global median per task (1141 ms for sentences; 801 ms for lists; 758 ms for words). Specifically, for each task we first excluded trials with outlier response times, which we defined as those in the bottom 2.5% or top 5% per participant. We then calculated median response times per task across participants (1,142 ms for sentences, 801 ms for lists, and 758 ms for words), and for each electrode and each trial, concatenated (a) the stimulus-locked data from 150 ms post-stimulus to 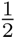 the median response time with (b) the production-locked data from 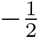 median response time to 150 ms pre-speech. We then linearly interpolated this time series to the number of time samples that would, when concatenated between the 150 ms post stimulus (stimulus-locked) and 150 ms pre-speech (speech-locked), result in a time series with the median planning period duration. Finally, we concatenated (a) the unwarped data leading up to 150 ms post-stimulus, (b) the warped data from the previous step, and (c) the unwarped data starting 150 ms before speech onset, forming the final epochs used in the analyses. We direct the reader to Morgan et al. 2024 for a fuller discussion and demonstration that this temporal warping increased signal to noise ratio in this dataset, as well as analyses of the unwarped high gamma activity across regions of interest.

## Acknowledgments

We thank the entire Flinker Lab for feedback on this project, and Yasamin Esmaeili for the Farsi example in the text. This work was supported by National Institutes of Health grants F32DC019533 (A.M.), R01NS109367 (A.F.), R01NS115929 (A.F.), and R01DC018805 (A.F.).

## Data and Code availability

Data will be made available from the authors upon request, provided documentation that the data will be strictly used for research purposes and will comply with the terms of our study IRB. The code is available at https://github.com/flinkerlab/decoding_words_in_sentences.

## Declarations

The authors declare that they have no competing interests.

## Appendix A Supplementary Information

### A.1 Stimulus Noun Properties

**Fig. S1.**
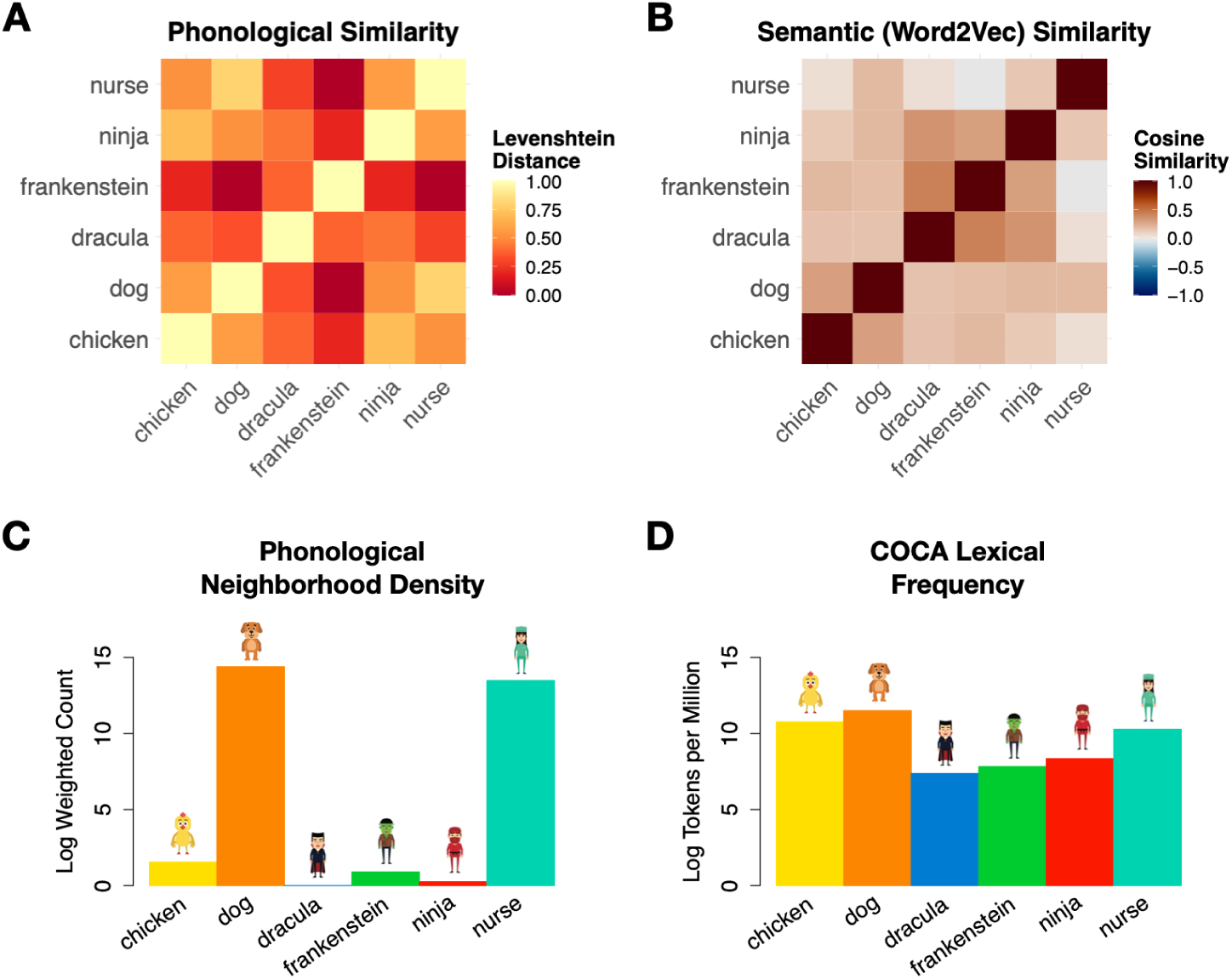
Psychologically relevant properties of the 6 nouns used in the stimuli. (A) Phonological similarity (scaled Levenshtein distance between phonological transcriptions). (B) Semantic similarity (Word2Vec embeddings). (C) Phonological neighborhood density (unstressed), weighted by the Kucera-Francis log frequency of each word’s neighbors. (D) Lexical Frequency taken from the Corpus of Contemporary American English (COCA) Version 2.0 (lemma instances per million words, logged).

### A.2 Data Warping

Supplementary Fig. S2, reproduced from Morgan et al. 2024, demonstrates the temporal warping procedure for a sample electrode (see Methods Sec. 4.8).

**Fig. S2.**
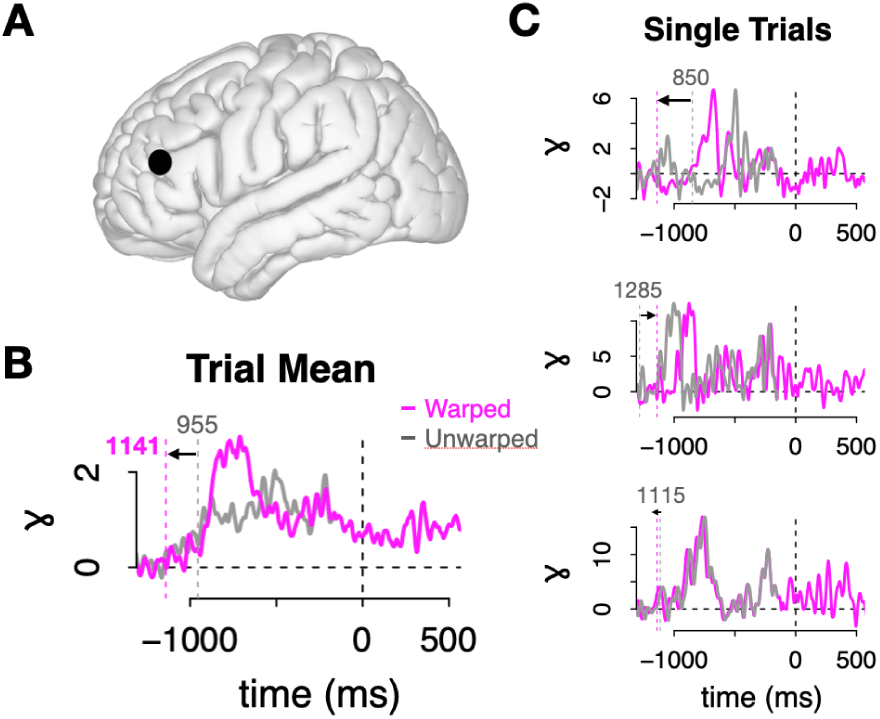
Warped and unwarped sentence production data from a sample electrode. The data in each trial between 150 ms post stimulus and 150 ms pre-speech were linearly interpolated to set the duration of the planning period to the global median per task (1142 ms for sentence production) [66]. (A) Sample electrode localization in MFG. (B) The mean of this electrode’s warped (pink) and unwarped (gray) trials. Prior to warping, this patient’s median sentence response time was 995 ms; after warping it was 1,141 ms: the median sentence production response time across patients. The peak of the warped data was higher than the unwarped peak, a sign that warping resulted in better temporal alignment and consequently higher signal-to-noise ratio. (C) Three sample trials: warped (pink) and unwarped (gray) data.

### A.3 Word-specific networks

**Fig. S3.**
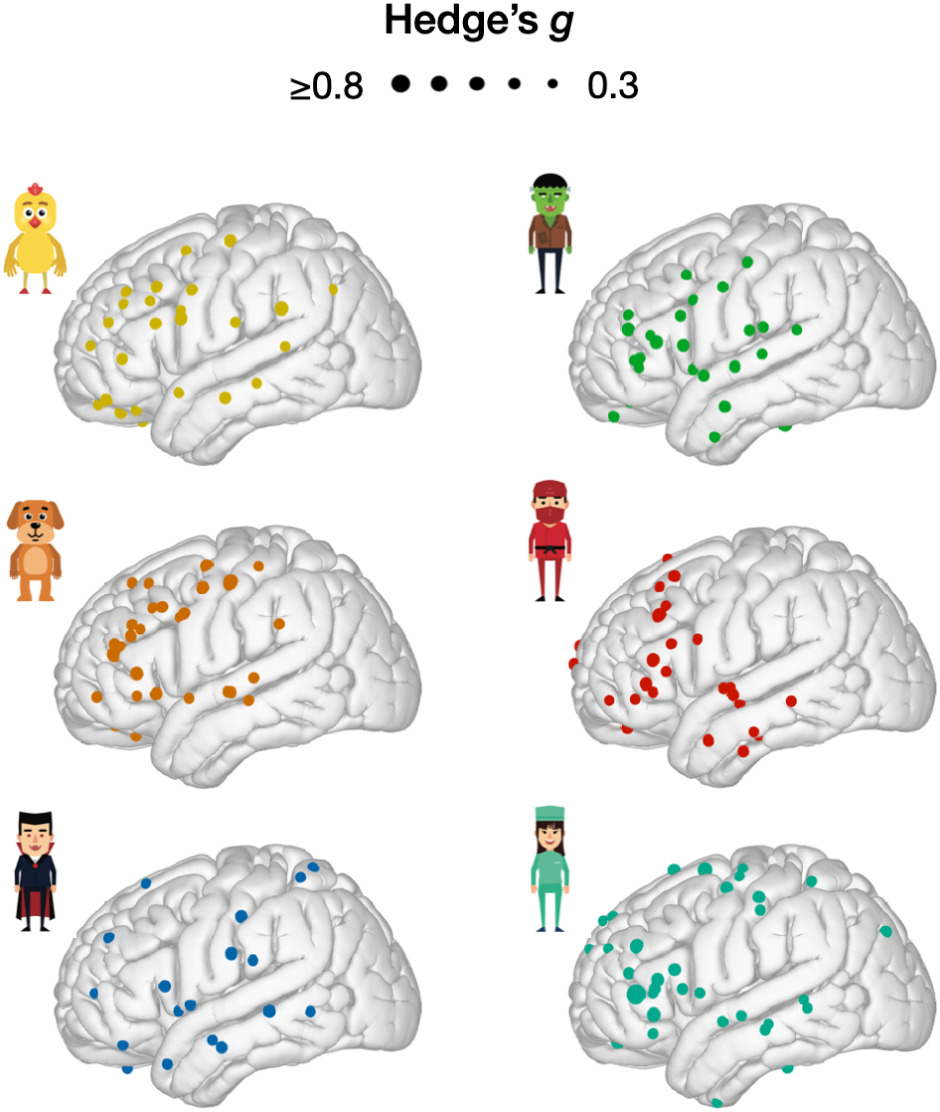
Cortical networks for each word. Each electrode’s activity for each word during picture naming was compared to that of all other words in each of the twenty consecutive 50 ms windows starting at -750 ms from speech onset. Electrode sizes correspond to the maximum difference across time, standardized by the pooled variance (Hedge’s *g*, a standardized difference metric interpreted like Cohen’s D). Only showing electrodes for which only one word has a maximum Hedge’s *g* exceeding 0.3 (i.e., just electrodes that uniquely encode one character).

### A.4 Picture naming classification results by ROI

**Fig. S4.**
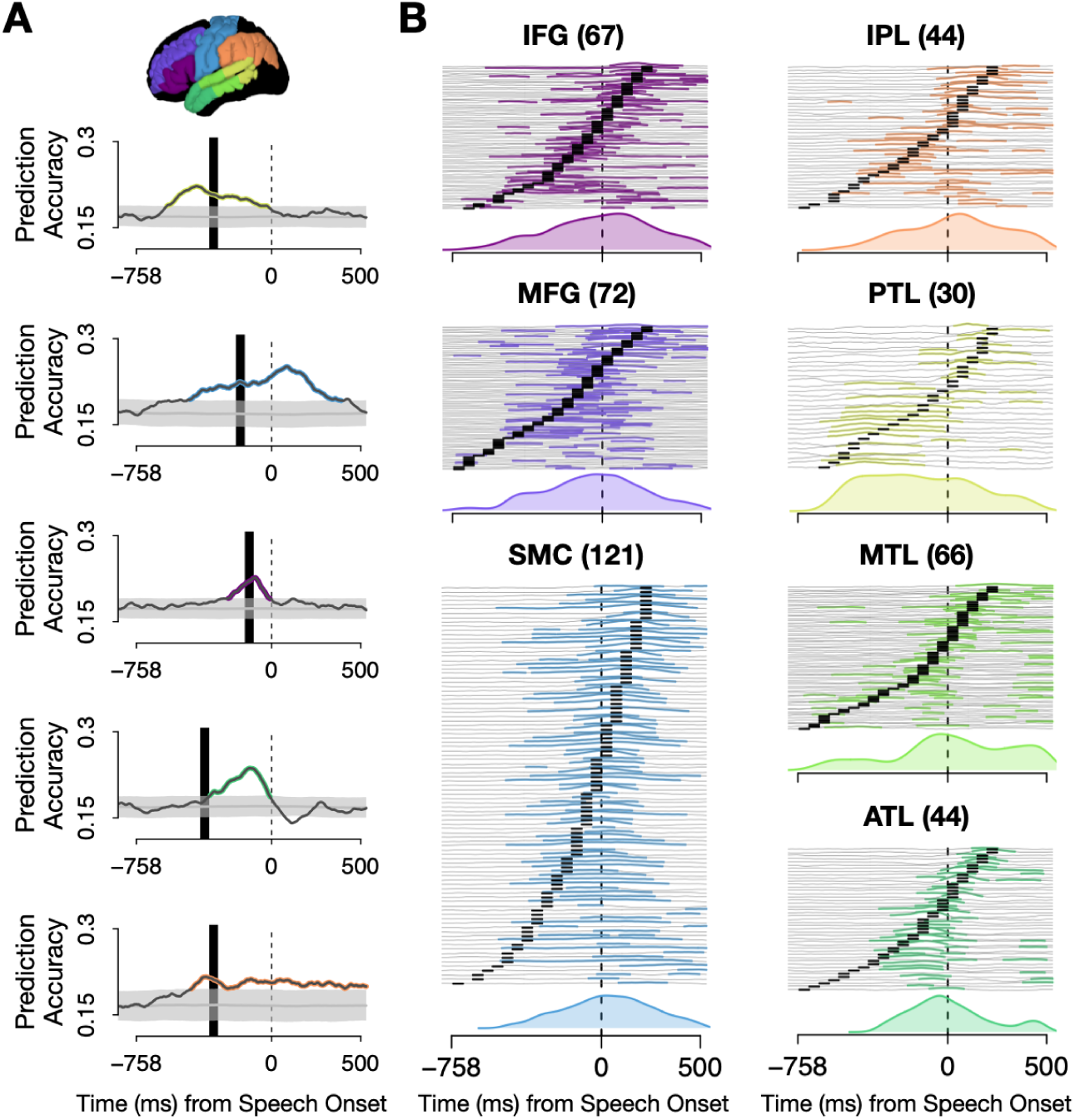
Picture naming classification results by ROI. (A) Results from five sample classifiers (see Fig. 2 for methods). From top to bottom, the classifiers were trained and tested on picture naming data in (1) Patient 9, PTL, -350 to -300 ms; (2) Patient 6, SMC, -200 to -150 ms; (3) Patient 9, IFG, -150 to -100 ms; (4) Patient 10, ATL, -400 to -350 ms; and (5) Patient 4, IPL, -350 to -300 ms. (B) Stacked picture naming prediction accuracies from each significant classifier by ROI (across patients and training windows). Number of significant classifiers appears in parentheses. Word identity was decodable in all seven ROIs, with the highest decodability in SMC and lowest in PTL, which was also the only ROI to peak during the pre-articulatory period.

### A.5 Sentence classification results by ROI

**Fig. S5.**
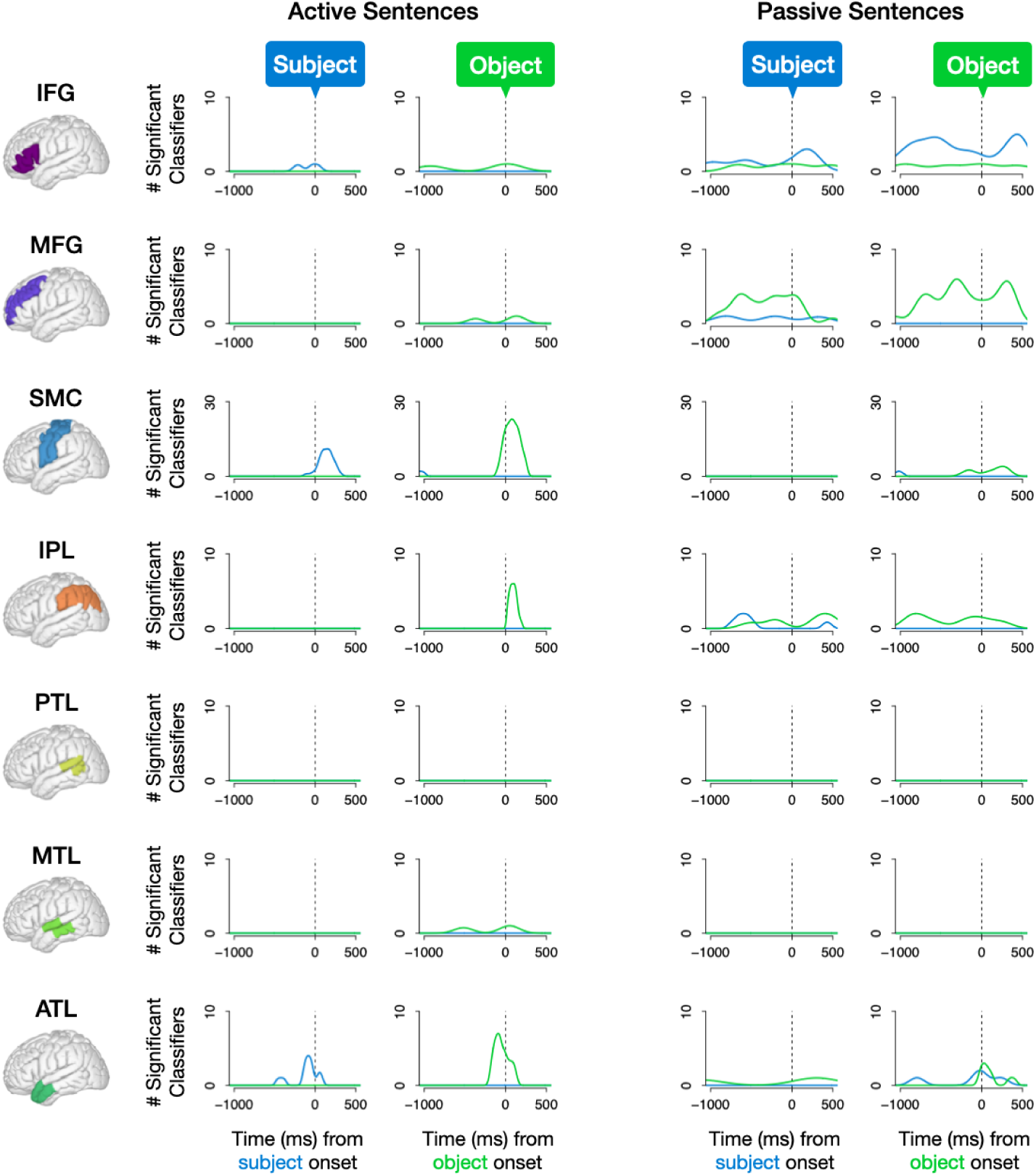
Sentence production classification results by ROI: Density of significant prediction accuracies of subject and object in active (left) and passive (right) sentences. The y-axis is scaled to the number of significant classifiers in each region and for both syntactic structures. (NB: Due to high decodability in SMC its vertical axis was scaled to 30; all other ROIs range from 0 to 10.) Notably, incongruent detections (i.e., predicting the subject during production of the object or predicting the object during production of the subject), which nearly exclusively occurred in passive sentences (Fig. 4E), appear to have been driven by processing in IFG, which preferentially encoded passive subjects, and MFG, which preferentially encoded passive objects.

### A.6 Reaction times

Median reaction times per task were 758 ms for picture naming; 801 ms for list production, 1,164 ms for active sentence production, and 1,424 ms for passive sentence production. Patients took significantly longer to begin passive sentences than both active sentences (*W* = 20493, *p <* .001, Wilcoxon rank sum test) and lists (801 ms; *W* = 9705, *p <* .001, Wilcoxon rank sum test).

## Footnotes

1 We elicited actives and passives by priming patients with questions in either active or passive syntax. This successfully elicited active and passive utterances from patients, although, consistent with previous literature [60], active priming was more successful in eliciting actives (96.93% of trials) than passive priming was in eliciting passives (49.15%). A mixed-effects logistic regression modeling passive production as a function of prime syntax converged with a random intercept for patient revealed that passive primes did indeed result in more passive responses than active primes (*β* = 4.242, *z* = 9.506, *p <* .001).

2 The Test of Equal Proportions was chosen due to the bounded nature of the counts, which were between 0 and 1280 (i.e., the total number of classifiers). Pairwise tests were only performed where the number of detections was higher for passives than actives because the reverse pattern is uninterpretabl*_√_*e: active analyses having roughly three times more data than passives (see Footnote 1), statistical power was 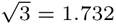 times higher for actives. The higher number of active detections is therefore likely trivial. On the other hand, the significantly higher number of incongruent detections for passives than actives, in spite of passives’ lower power, lends extra credibility to those effects.

## Notes

https://github.com/flinkerlab/decoding_words_in_sentences

### Competing Interest Statement

The authors have declared no competing interest.

### Summary of Updates

This version offers longer discussions of previous models of word production, and shifts the focus of the findings to emphasize (1) an interpretation of our findings in prefrontal cortex as a mechanism for top-down control of sentence production and (2) the encoding of syntactic roles -- specifically, the representation of subjects in IFG and objects in MFG.

